# Unraveling Keystone Taxa: Interactions Within Microbial Networks and Environmental Dynamics in Lake Mendota

**DOI:** 10.1101/2024.11.11.623027

**Authors:** Qiyao Yang, Rosa Aghdam, Patricia Q. Tran, Karthik Anantharaman, Claudia Solís-Lemus

## Abstract

Microbial communities in freshwater ecosystems drive critical biogeochemical cycles, nutrient transformations, and energy flows essential for ecosystem stability. Yet, in the face of accelerating environmental changes, the responses of these microbial networks to spatial and temporal shifts remain underexplored, particularly with rising anoxia. We investigated the microbial ecosystems of Lake Mendota, Wisconsin, USA, through comprehensive metagenomic and metatranscriptomic analyses to elucidate their adaptations to environmental fluctuations across temporal and spatial dimensions. Employing tools like Sparse Inverse Covariance Estimation for Ecological Association Inference (`SPIEC-EASI`) and Conditional Auto-Regressive Least Absolute Shrinkage and Selection Operator (`CARlasso`), we identified key microbial taxa and their interactions with environmental parameters such as depth, temperature, pH, and dissolved oxygen. Our findings reveal that biological interactions more than environmental variables shape microbial community assembly and function. Specifically, keystone taxa from the phylum *Bacteroidota* emerged as pivotal in nutrient cycling and organic matter decomposition, processes crucial for sustaining water quality. Notably, these keystone taxa demonstrate dynamic adaptability, suggesting that microbial networks can rapidly adjust to changes in composition, a trait essential for resilience in the face of warming temperatures and altered precipitation patterns. This study provides critical insights into the resilience and adaptability of freshwater microbiomes, highlighting the role of microbial interactions in maintaining ecosystem health. By understanding how these microbial networks respond to environmental pressures, we can better predict shifts in microbial dynamics and anticipate the broader ecological impacts of climate change on freshwater systems.

**Importance:** This research underscores the critical role of keystone taxa in freshwater ecosystems, highlighting how these organisms maintain water quality and contribute to the stability of aquatic environments. Understanding the ecological roles of these taxa is essential for developing strategies to manage ecosystems and conserve freshwater resources, particularly in the face of ongoing environmental challenges like climate change. The insights provided by this study not only enhance our comprehension of microbial interactions but also support effective ecosystem management and conservation efforts.

## 1 Introduction

The study of microbial communities within aquatic ecosystems offers critical insights into ecological processes that maintain environmental health and biodiversity [1]. These communities are essential players in biogeochemical cycles [2], contributing to nutrient transformation [3], energy flow [4], and the overall stability of ecosystems [5]. With rapid advancements in molecular biology and computational analysis, the field of microbial ecology has greatly expanded our understanding of how microorganisms interact with their environment and each other under various ecological settings [6, 7].

Aquatic ecosystems, particularly those in freshwater environments like lakes, present a dynamic environment where microbial populations are subject to a variety of physical, chemical, and biological forces. Environmental factors such as water temperature [8], dissolved oxygen level [9], and the presence of dissolved organic matter [10] create a highly heterogeneous habitat that influences microbial diversity and function. Understanding these intricate microbial dynamics is essential not only for advancing ecological research but also for informing effective environmental management and conservation strategies, thus ensuring the resilience and sustainability of these vital ecosystems.

Recent advancements in molecular biology and computational analysis have significantly enhanced our understanding of microbial communities in aquatic ecosystems. These studies underscore the importance of physical, chemical, and biological factors in shaping the structure and function of microbial populations. For instance, variations in salinity, temperature, and nutrient availability have been found to profoundly influence microbial diversity and dynamics in both saline and freshwater environments [11] [12]. This body of research, utilizing sophisticated statistical models, has not only elucidated the complex interactions within microbial communities but also emphasized their pivotal role in maintaining ecosystem health and stability. Bryanskaya et al. [11] specifically explored the microbial communities of saline lakes in the Novosibirsk region, revealing that salinity and other geochemical parameters significantly affect microbial composition and abundance. Their use of fluorescent in situ hybridization (FISH) provided a direct visualization of microbial communities, demonstrating the dominance of Archaea in higher salinity conditions. This study exemplifies how environmental variables like salinity can shape microbial landscapes, thus offering insights crucial for ecological research and effective conservation strategies. Such insights are essential for developing strategies for conservation and understanding the ecological impacts of environmental changes on microbial populations in aquatic systems.

Similarly, in the realm of soil microbiology, a study by Banerjee et al. [13] provides critical insights into the complex interactions within microbial ecosystems. Utilizing sequencing alongside sophisticated network analysis, this research dissects the co-occurrence patterns of bacterial and fungal communities in response to organic matter decomposition. The application of network analysis has illuminated the dynamics between microbial taxa and environmental variables, particularly how nutrient additions reshape these interactions. Notably, the study identifies key taxa such as Acidobacteria, Frateuria, Gemmatimonas, Chaetomium, Cephalotheca, and Fusarium, which exhibit strong positive correlations with decomposition processes. These keystone taxa are pivotal in nutrient cycling and demonstrate a substantial influence on the ecological functionalities of their respective communities. The methodology and findings in [13] offer a valuable perspective on the functional redundancies within microbial communities, stressing the importance of keystone species in maintaining ecological balance and process efficiency. Further expanding on this topic, a more recent study by Zhang et al. [14] examines how microbial assemblages within different soil aggregate sizes impact nutrient cycling in agroecosystems, highlighting the influence of both deterministic and stochastic processes on these communities and their functional roles in carbon, nitrogen, and phosphorus cycling.

We applied this framework to identify keystone species which might disproportionally influence the ecosystem in a model, dynamic lake ecosystem (Lake Mendota, WI, USA). Building upon a prior dataset from this lake [15], we reanalyzed the data with this new perspective. In summary, the water column of Lake Mendota (WI, USA), was previously sampled weekly and at various depths (5, 15, 20, and 23.5m) over the course of the summer and early fall of 2020. These paired metagenome and metatranscriptome samples span dynamic environmental conditions. These conditions are primarily summer anoxia, during which half of the water column lacks oxygen, and water column mixing, which occurred on October 18, 2020. During that sampling, water was collected for both metagenomics (bulk DNA) and metatranscriptomics (RNA) to determine microbial composition, abundance, and activity. The data was previously binned into a set of 431 metagenome-assembled genomes (a.k.a MAGs) (see [15] for detailed methodologies). Here, our investigation focuses on how microbial communities adapt to temporal and spatial variations, aiming to unravel the intricate network of interactions that maintain microbial diversity and functionality [16, 17, 18]. Employing a suite of statistical tools and visual methods, our study illustrates the distribution and variability of microorganisms across spatial and temporal gradients (i.e., different sampling times and depths). We provide key insights into the relationships among microbial taxa, revealing underlying ecological interactions crucial for understanding community dynamics. We further explore the intricate relationships between MAGs and various environmental parameters such as depth, temperature, pH, and dissolved oxygen [19, 20, 21]. By using the Sparse Inverse Covariance Estimation for Ecological Association Inference (`SPIEC-EASI`) [22] and the Chain Graph Least Absolute Shrinkage and Selection Operator (`CARlasso`) [23, 24], our research identifies key MAGs and constructs a detailed Gaussian graphical model to discern significant relationships between microorganisms and environmental conditions. The findings highlight key microbial taxa with significant connections to environmental factors, underscoring their adaptive or functional roles within the lake ecosystem [25, 26].

## 2 Results

### 2.1 Comparative and Correlative Analysis of Microbial Abundance in Varied Environmental Conditions

In this study, we analyze 16 samples collected from Lake Mendota during the summer and early fall of 2020 to investigate microbial dynamics across different environmental conditions. These samples were examined at multiple depths and included both metagenome and metatranscriptome data, allowing for a comprehensive analysis of microbial community structure and function. Our analysis centers on 431 metagenome-assembled genomes (MAGs), employing various statistical tools to understand the intricate relationships between microbial taxa. We first analyzed the microbial abundance patterns in the 16 samples. This analysis revealed significant temporal and spatial variations in microbial community dynamics, with (Figure S1) pre-normalization analyses showing notable variability in abundance ranges across different samples, and post-normalization analyses (Supplementary Figure S2 providing a clearer view of these variations by illustrating the central tendency and outliers more distinctly. The dataset was categorized by collection month—July (3 samples), August (6 samples), September (2 samples), and October (5 samples)—to facilitate the exploration of the microbial community’s temporal dynamics. Our temporal analyses (Figure 1a) highlighted that August, which had the highest number of samples, also exhibited more outliers compared to other months. This occurrence of outliers is likely a consequence of the larger sample size in August, indicating significant deviations in relative abundance from the monthly trends. Ecologically, as the summer progresses, the lake becomes increasingly stratified, leading to a clearer distinction between the top and bottom layers—characterized by differences in physico-chemical conditions such as temperature, light, oxygen, etc.— which could influence species richness and diversity. Additionally, a binary heatmap was constructed for the top ten most prevalent genomes, illustrating their presence across the months (Figure 2). This visualization outlines the temporal patterns and dominance of these key genomes, providing valuable insights for environmental monitoring and microbiome research. Of the MAGs analyzed, 29 belonging to 8 different phyla were ranked among the top ten in terms of abundance across the dataset. Some MAGs, for example, members of Actinobacteria, were only ranked in the top 10 in October, whereas members of Verrucomicrobia were ranked in the top only before October. Other phyla, such as Proteobacteria, showed more diverse patterns (some were only in the top 10 before or during October) reflecting the broad ecological niches that this phylum occupies.

**Figure 1.**
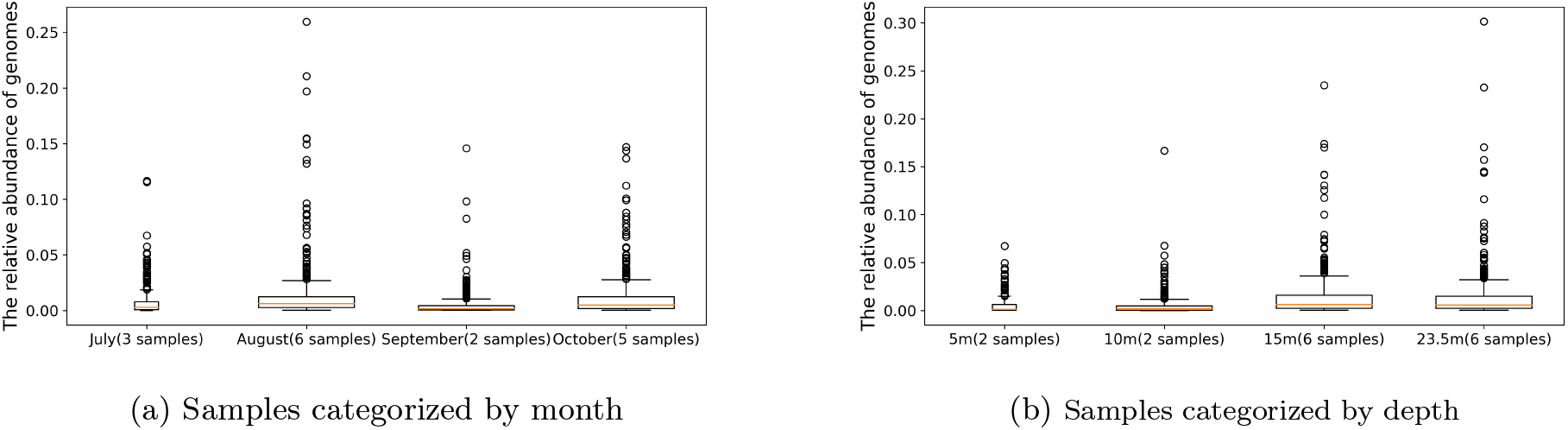
Sum of the relative abundance of each genome based on the normalized dataset (row sum normalization method): (a) categorized by month, and (b) categorized by depth.

**Figure 2.**
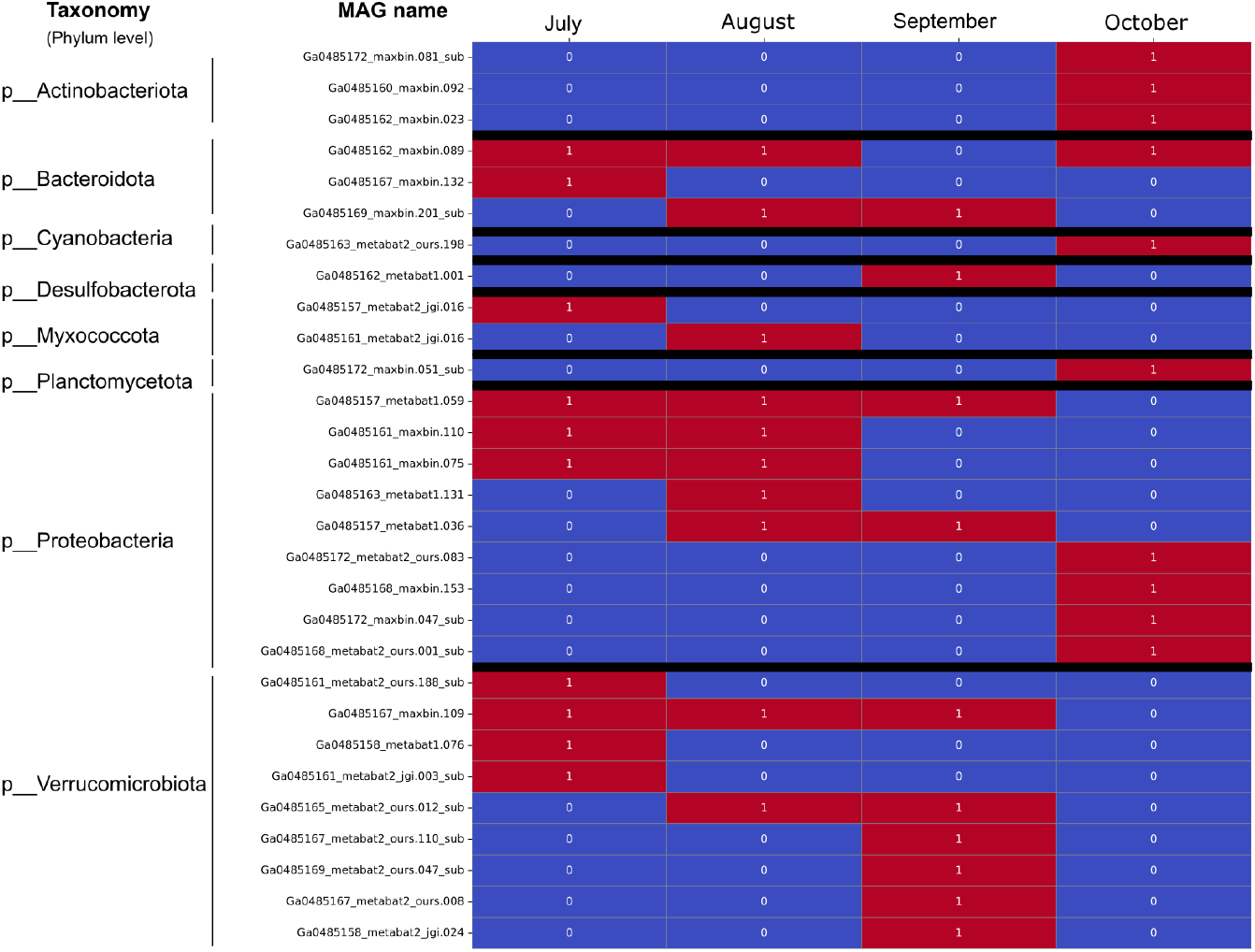
Binary heatmap showing the occurrence of the top ten most prevalent genomes for each month. The rows represent different MAG names, categorized by their taxonomy at the Phylum level. The colors indicate the presence (red) and absence (blue) of genomes in the top ten most popular genomes for each month.

Water column depth is associated with decreased oxygen concentrations during the stratified period, along with distinct biogeochemical profiles. To study the impact of water column depth on the microbial community, we analyzed samples originating from different depths: 5 meters (2 samples), 10 meters (2 samples), 15 meters (6 samples), and 23.5 meters (6 samples). We observed distinct patterns in genome abundances at each depth (Figure 1b). Notably, deeper samples showed a higher occurrence of outliers which was expected given the fewer samples at 5 and 10 meters, potentially influencing observed counts. We identified central tendencies, variability, and outliers, thereby elucidating the abundance dynamics specific to each depth. We also identified the top ten most prevalent genomes at each depth (Figure 3). In total,30 MAGs from 8 distinct phyla ranked among the top ten in terms of abundance. For instance, MAGs from the phylum Planctomycetota appeared in the top ten exclusively at a depth of 5m, while Bacteroidota MAGs only ranked in the top ten at depths beyond 5m (i.e., at 10m, 15m, and 23.5m). Proteobacteria displayed a more varied distribution, with some MAGs only appearing in the top rankings beyond the depth of 10m and showing less presence at 5m, which illustrates the extensive ecological diversity this phylum exhibits across different water depths.

**Figure 3.**
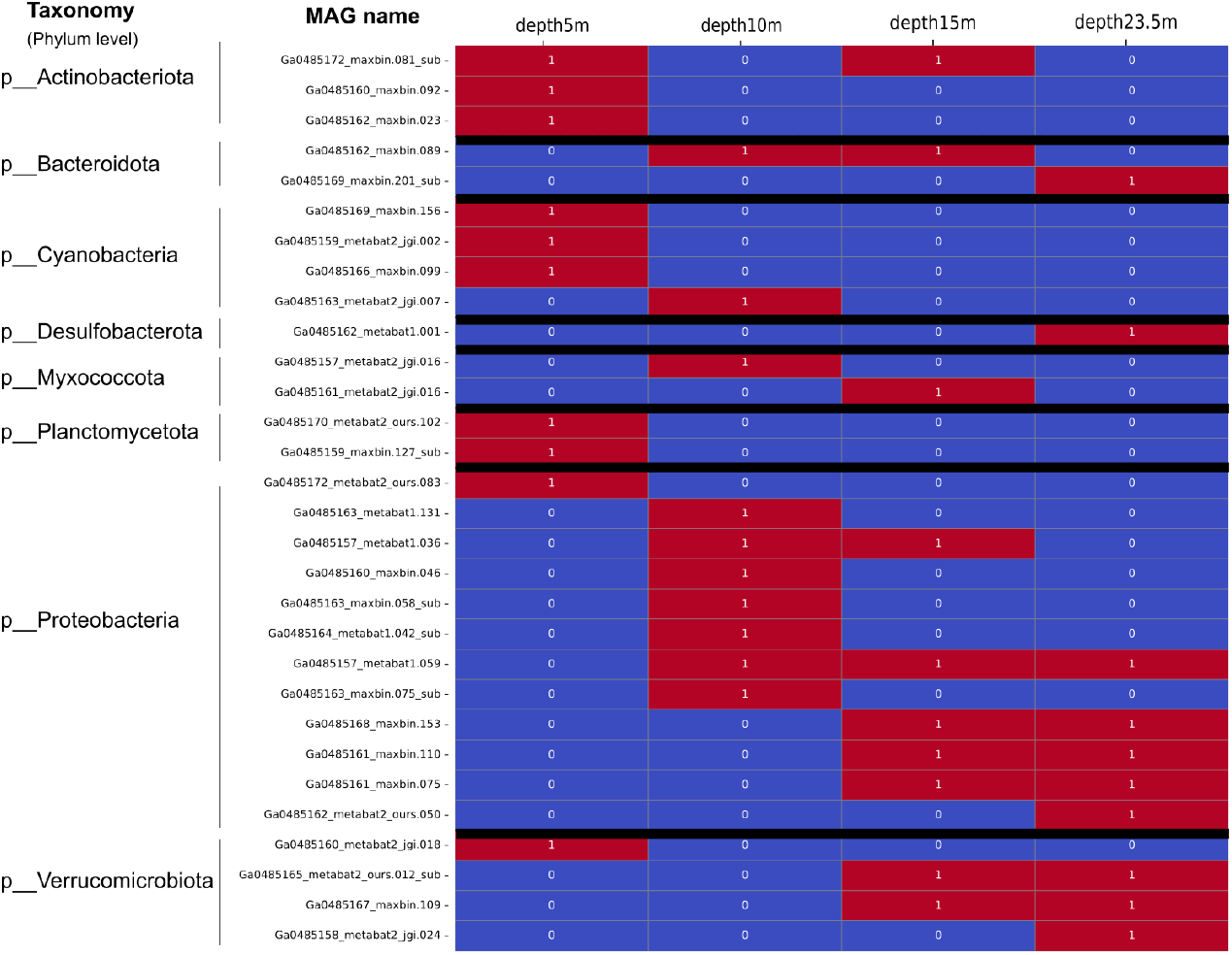
Binary heatmap for the occurrence of the top ten most popular genomes for each depth. The rows represent different MAG names, categorized by their taxonomy at the Phylum level. The colors indicate the presence (red) and absence (blue) of genomes in the top ten most popular genomes for each month.

To delve deeper into the microbial community dynamics within the ecosystem, we conducted a comparative analysis of the relative abundance of the top ten most prevalent genomes across different time periods (July, August, September, and October) and depths (5 meters, 10 meters, 15 meters, and 23.5 meters) (See Figure S3). We studied patterns in genome abundance, which can aid in understanding the dynamics of microbial communities over time and depth, offering a nuanced view of how microbial community composition shifts over time and with depth. This comparative analysis not only highlights the shared presence or exclusivity of key microorganisms but also sets the stage for exploring the complex interactions between microbial communities and environmental factors, enhancing our understanding of how temporal and spatial factors influence microbial dynamics.

Next, to assess interrelationships and co-occurrence patterns among the 16 samples, we performed correlation analyses using both Pearson’s and Spearman’s methods, which yielded consistent correlation values across the samples, thereby confirming robust associations regardless of the method used. This consistency underscores the reliability of our findings and is visually summarized in Supplementary Figures S4 for Pearson correlation and S5 for Spearman correlation. These results suggest that both linear and monotonic associations are prevalent in the microbial community, providing a comprehensive understanding of microbial interactions within the ecosystem. Notably, the strong correlations observed between samples collected from different depths on the same date highlight the influence of depth and temporal factors on microbial community structure.

### 2.2 Using CARlasso to Identify Connections Between MAGs and Environmental Parameters

#### 2.2.1 Identifying Central MAGs in Microbial Community Networks

Building from our analyses of microbial community dynamics, we used raw count data from bulk metagenomic sequencing to explore connections between bacterial and archaeal MAGs and various environmental parameters. Using `SPIEC-EASI`, followed by degree calculation, we identified central MAGs in the community and constructed a Bayesian sparse microbial network via the `CARlasso` method [23, 24] to investigate interactions among microbes and environmental influences on microbial composition. Our approach revealed how specific MAGs correlated with environmental variables such as pH, temperature, dissolved oxygen, chlorophyll, etc.

Our analysis focused on the “top 15 nodes” resulting by maximum degree in the microbial network (Section 4.2.1) with all environmental features within a microbial network (Figure 4). Degree centrality measures the number of direct connections, or edges, that a node has. In this context, we identified and examined the MAGs that exhibited the highest degree of connectivity, reflecting their active engagement with other microbes and environmental parameters. Such nodes are considered highly interactive within the network and potentially play crucial roles in microbial processes and environmental interactions. Overall, this analysis uncovered the most connected taxa which may have a significant impact on the structure and function of the ecological community. Detailed node identities and their corresponding phylum level associations are provided in Table 1.

**Table 1:**
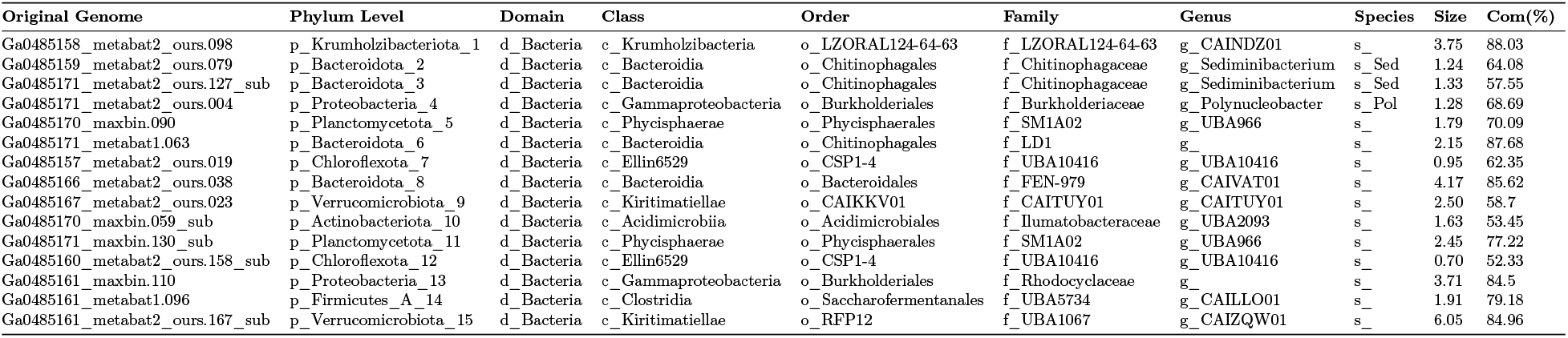
Description of the names of the nodes in this network: Original Genome to Completeness. This table serves as a reference for the response nodes in the above network. The “Original Genome” column represents the MAGs’ names, while the other columns indicate the corresponding taxonomic and genome details for each MAG as depicted in the network. In the Species column, s_Sed refers to s_Sediminibacterium sp002299885 and s_Pol refers to s_Polynucleobacter sp002292975. The “Size” column represents genome size in megabase pairs (Mbp), and the “Com(%)” column indicates completeness percentage.

**Figure 4.**
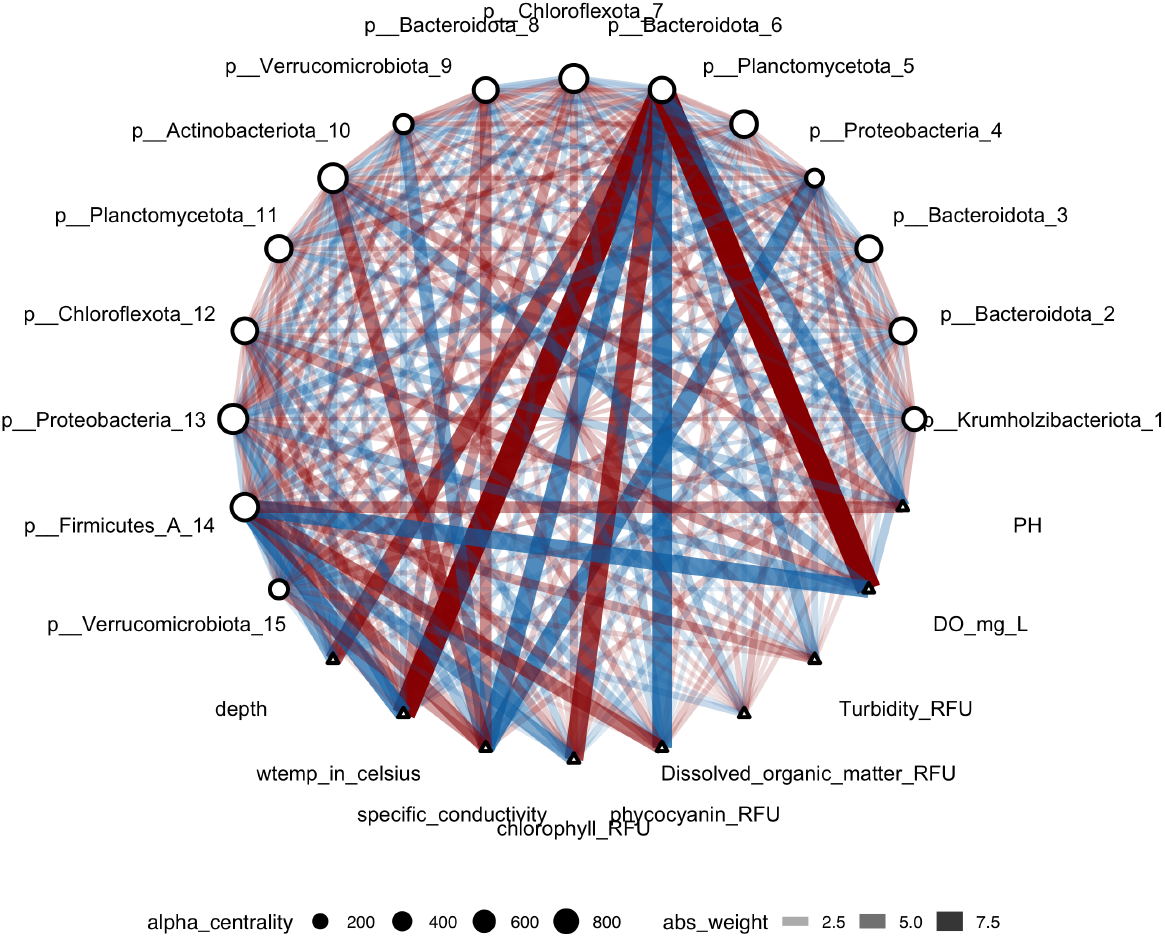
`CARlasso` network diagram showing the top 15 nodes with the highest degree centrality, selected using the `SPIEC-EASI` algorithm [22], and their connections to environmental features. The nodes, representing MAGs categorized by Phylum, and environmental parameters, are connected based on the `CARlasso` analysis [23, 24]. Node size indicates degree centrality and edge thickness reflects interaction strength. The edge color indicates the type of relationship, with blue representing negative relations and red representing positive relations. This visualization highlights key microbial taxa and their interactions with environmental variables like pH, water temperature, and dissolved oxygen, illustrating their significant ecological roles.

#### 2.2.2 Assessing the Impact of Community Changes

Focusing centrality of this MAG Bacteroidota_6 (Ga0485171_metabat1.063), we sought to study the role of other members in influencing its “key” status of Ga0485171_metabat1.063. To do this, we performed a permutation analysis (n=100 times) where all other MAGs in the previous analysis, except for Bacteroidota_6, were swapped with 14 other randomly selected MAGs. This process aimed to examine the impact of Bacteroidota_6 on the network’s structure and functionality by observing how the network adapts to the reassignment of this highly connected genome. We observed a significant change with its previously strong connections with various environmental features dissipating in these analyses (Figure 5). This shift highlights the community-dependent nature of Bacteroidota_6’s role and underscores the dynamic adaptability of microbial networks to changes in their composition. A comprehensive list of changes in the network nodes is provided in Table 2, which lists each node’s original genome name and its corresponding Phylum, providing essential context for interpreting the impact of the permutations on the network. Interestingly, when permutations revealed the loss of centrality of Bacteroidota_6, some alternative members of the community gained key importance. For example, we observed a three-species interaction between members of Actinobacteria, Bacteroidota, and Cyanobacteria.

**Table 2:**
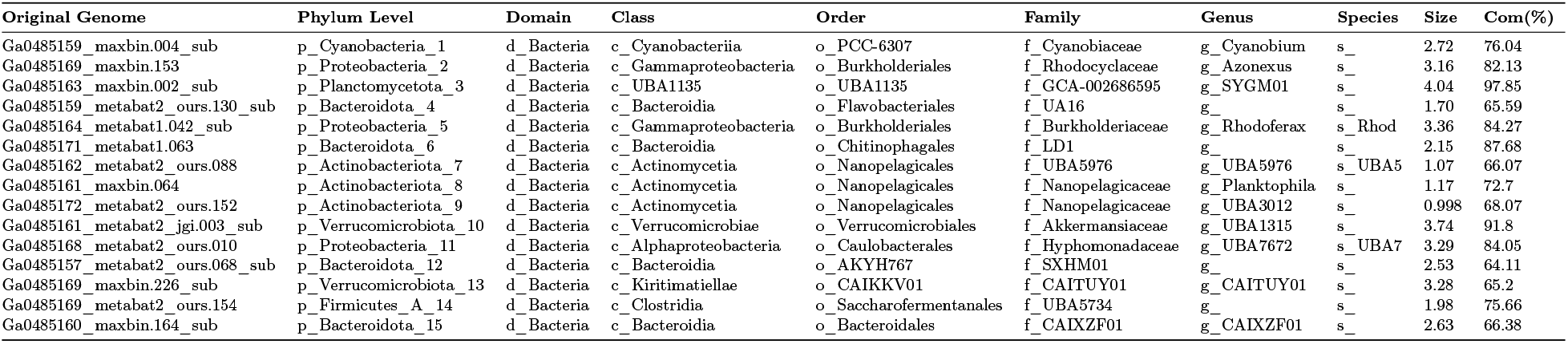
Description of the names of the nodes in this network: Original Genome to Completeness. This table serves as a reference for the response nodes in the above network. The “Original Genome” column represents the MAGs’ names, while the other columns indicate the corresponding taxonomic and genome details for each MAG as depicted in the network. The “Size” column represents genome size in megabase pairs (Mbp), and the “Com(%)” column indicates completeness percentage. s_ Rhodoferax sp007280205 refers to s_ Rhod, s_UBA7672 sp903842595 refers to s_UBA7, and s_UBA5976 sp903951185 refers to s_UBA5.

**Figure 5.**
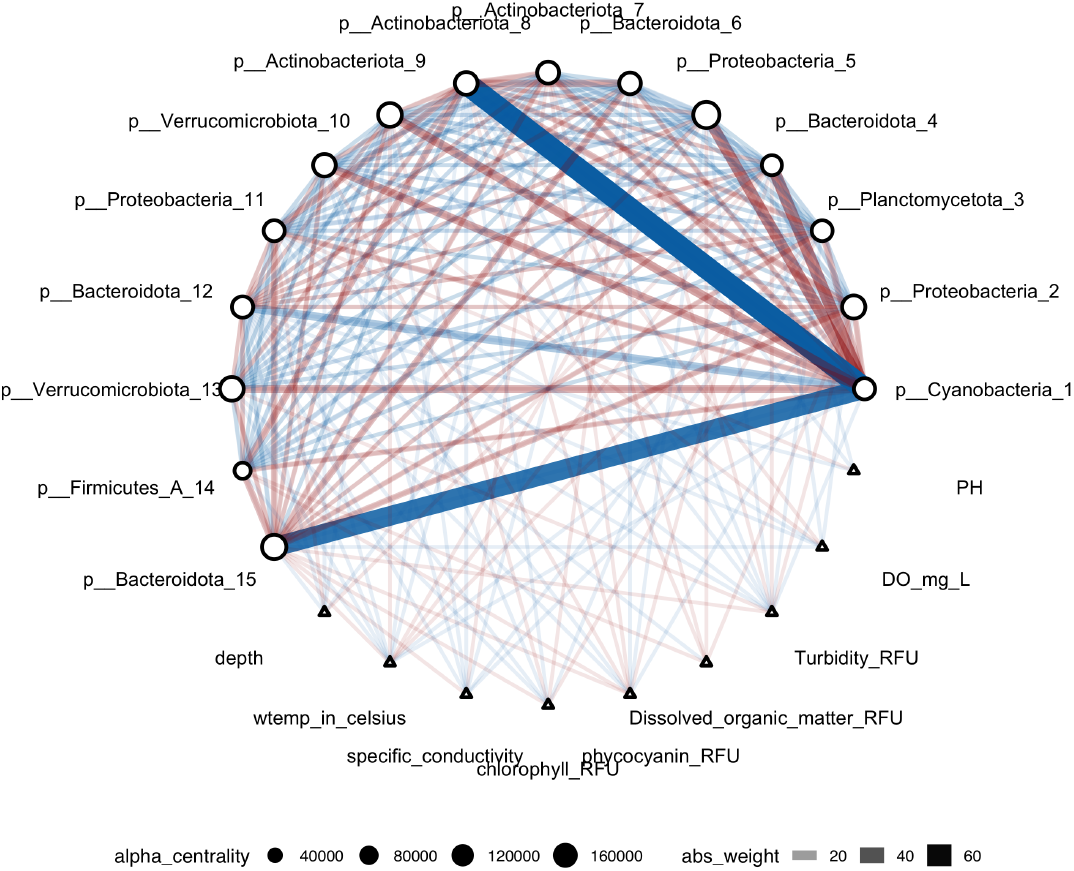
Permutation on p_Bacteroidota_6 with All Environmental Features. This network diagram shows the results of the permutation analysis, where p_Bacteroidota_6 was swapped with 14 randomly selected MAGs. Nodes represent MAGs and environmental parameters, with edge thickness indicating interaction strength (red for positive, blue for negative). Node size reflects degree centrality. The analysis highlights the significant role of p_Bacteroidota_6 in the network, with notable changes in its connections to environmental features.

#### 2.2.3 Comparative Network Analysis of Key Connectors Within Simulated Microbial Community Networks

Next, we posited that the presence of taxa in the environment alone does not necessarily indicate that they are active. To further designate a MAG as a key taxon, we proposed that the microorganisms must actively interact with the environment and leverage gene expressions from metatranscriptome data to answer this question. We conducted a detailed analysis of two microbial genome-assembled genomes (MAGs), Ga0485171_metabat1.063 and Ga0485167_metabat2_ours.023, comparing their roles within metagenomic and metatranscriptomic contexts across simulated microbial community networks. Using permutation tests and analyzing conditional regression coefficients via the `CARlasso` model [23, 24], we assessed the connectivity and ecological importance of these MAGs in different metatranscriptomic (activity) network configurations. In our comparative analysis of microbial community dynamics, we randomly created a microbial community with Ga0485171_metabat1.063 (Bacteroidota_6) and 14 other taxa, with 100 repetitions. During each permutation, we tracked when any edge values exceeded the 90th percentile in the distribution of all edge values – an indicator of the MAG’s connectivity. These edge values represent conditional regression coefficients generated by the `CARlasso` model. This criterion was met in 42% of metagenomic permutations and 44% of metatranscriptomic permutations. The detailed results are documented in Table 3, which illustrates the significant role this MAG plays across diverse environmental settings, and its slightly higher significance in gene expression activities compared to genomic presence. We conducted similar analyses for another MAG, Ga0485167_metabat2_ours.023 (Verrucomicrobiota_9). Our results 3 indicated that Ga0485167_metabat2_ours.023 met this significance threshold in only 11% of metagenomic permutations and 6% of metatranscriptomic permutations. This suggests that while this MAG is present in the microbial community, its role is markedly less pronounced than that of Ga0485171_metabat1.063, especially in the context of active metabolic processes.

**Table 3:**
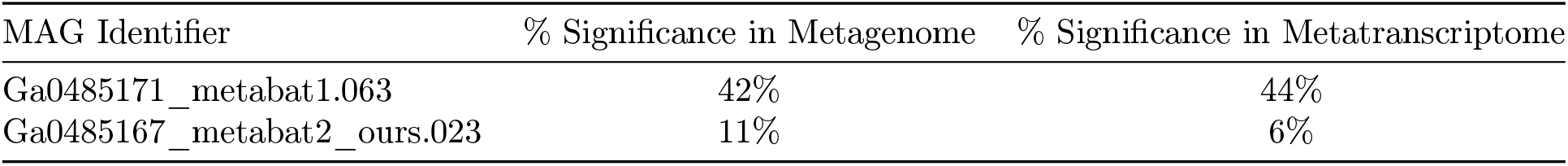
Comparative Significance of Microbial Genome-Assembled Genomes (MAGs) in Network Configurations. The table presents the percentage of significance for two MAGs, Ga0485171_metabat1.063 and Ga0485167_metabat2_ours.023, in both metagenomic and metatranscriptomic network configurations. Significance is determined by the percentage of permutations in which edge values exceeded the 90th percentile, indicating the MAG’s connectivity and ecological importance within the microbial community.

| MAG Identifier | % Significance in Metagenome | % Significance in Metatranscriptome |
| --- | --- | --- |
| Ga0485171_metabat1.063 | 42% | 44% |
| Ga0485167_metabat2_ours.023 | 11% | 6% |

## 3 Discussion

### Microbial community composition influences who is a keystone taxa

Our analyses underscore that the role of key microbes is highly dependent on the composition of the surrounding microbial community. Whenever we identified a “key taxa”, we tested whether it remained a key taxon upon mixing up the other members in the community. Two patterns emerged: either the microbe retained its importance, or it lost its significance. When it lost its importance, other community members sometimes emerged as important, indicating the critical nature of specific interactions within a consortium of microbes. This reveals that the importance of certain microbes can be highly context-dependent and influenced by their microbial neighbors. Moreover, our findings highlight that biological interactions between community members can sometimes outweigh environmental factors in determining the structure and functionality of microbial networks. This dynamic adaptability underscores the complexity of microbial ecosystems and suggests that the ecological roles of microbes are not fixed but can shift depending on community composition. Understanding these complex interactions is vital for comprehending broader ecosystem dynamics and the specific roles played by different microbes across various environmental contexts. Recent scholarly work, including a comprehensive review by Tudela et al. [27], has further explored the indispensable roles of keystone species within the gut microbiota—key regulators of metabolic functions and host health. These keystone species significantly influence the stability of microbiomes; their presence or absence can drastically alter microbiome architecture and, consequently, affect the host’s health profoundly. Adding to this, Xun et al. [28] focused on soil microbiomes, demonstrating how keystone taxa with specialized metabolic functions, such as nitrogen and phosphorus metabolism, enhance community stability. The findings reveal that these keystone taxa play a pivotal role in maintaining soil health and fertility. Moreover, recent research highlights the essential role of keystone taxa in the recovery of the human gut microbiome after antibiotic treatment, identifying specific bacteria that facilitate rapid ecological recovery and resilience [29]. This underscores the broader ecological principle that keystone taxa are not only crucial for maintaining balance and health in natural ecosystems but also play a fundamental role in human health, particularly in the resilience and stability of our internal microbial communities. The implications of this work suggest that enhancing or protecting these keystone taxa could be key to managing microbiome health post-disturbance, further emphasizing the interconnectedness of microbial roles across different environments and health contexts.

### Activity measurements provide additional context for the identification of keystone taxa

In this study, we build upon metagenomic characterization of a freshwater lake microbiome to incorporate metranscriptomics to unravel keystone taxa, microbial and environmental interactions. These methodologies come with their own set of challenges. Metagenomics, which involves sequencing bulk DNA from the environment, typically portrays the “genomic potential” of ecosystems. Going a step further, metagenomic binning, reconstructs microbial genomes, and can therefore link genomic function and taxonomy [15]. Most prior work on the identification of keystone species has focused on amplicon (16S rNA sequencing) or metagenomic analyses. However, combining these analyses with experimental or quantitative data increases the resolution of observations and further decodes community interactions. With this in mind, we applied network analysis methods to not only identify species and environmental interaction in a complex, dynamic system, but also validate these “hypothetical keystone species” through simulating mock communities, and targeting active taxa. Network analysis tools like `SPIEC-EASI` [22] and `CARlasso` [23, 24] rely heavily on statistical correlations, which do not necessarily imply causation. Therefore, the predicted interactions they generate often require further validation through direct experimental manipulations or longitudinal observational studies to confirm their ecological relevance and functional significance [30, 31]. The integration of activity measurements, particularly those derived from metatranscriptomic data, has profoundly enhanced our understanding of microbial community dynamics and keystone taxa identification. This approach not only confirms the presence of taxa within a community but crucially illuminates their active roles and interactions [32]. Our analysis reveals how taxa like members of *Bacteroidota* exhibit significant ecological functions in freshwater lakes that are sometimes only detectable through gene expression profiling, underscoring their roles as keystone species within different network configurations. Furthermore, our findings align with the emerging consensus in the field that the structural importance of a microbe, as suggested by metagenomic data, does not always correlate with its functional impact. This is exemplified by Bacteroidota_6 whose lower activity levels contrast with its genomic presence, highlighting the need for a nuanced approach to defining keystone taxa that considers both presence and activity. Such distinctions are critical, as discussed in recent influential studies [33, 34], which advocate for a dual perspective on microbial significance in ecosystems. An additional component that can be integrated with microbial metagenomics and metatranscriptomics is viromics [35], which can unravel factors underpinning observed dynamics and lead to a comprehensive understanding of microbiomes.

### Metabolic basis for interactions

Members of complex microbiomes benefit from interactions among members. Through their metabolisms, in other words, through microbes ingesting, transforming, and releasing compounds in the environment, other microbes can thrive. In our analyses, we identified two microorganisms in the anoxic zone of Lake Mendota MAGs, Ga0485171_metabat1.063 (Bacteroidota_6 d_Bacteria c_Bacteroidia o_Chitinophagales) and Ga0485167_metabat2_ours.023 (p_Verrucomicrobiota_9 d_Bacteria c_ Kiritimatiellae o_CAIKKV01) that were defined as key species due to their connectivity, abundance and activity in the ecosystem. These two microbes, belonging to two different phyla, have commonalities in terms of their metabolism. Looking into their genomes and their metabolic potential, both organisms encode genes for amino acid utilization, arsenate reduction, complex carbon degradation, in particular hexosaminidase for chitin degradation. Specifically, Ga0485167_metabat2_ours.023 encodes genes for FeFe and Nifehydrogenes and methane oxidation. Neither encoded genes for oxygen metabolisms, hinting at their lifestyle as anaerobic taxa.

### Implications for ecosystem management

The insights garnered from our study of the microbial ecology of Lake Mendota have profound implications for the management and conservation of this vital freshwater ecosystem. Understanding the roles and interactions of microbial communities not only enriches our ecological knowledge but also informs practical strategies for ecosystem management. Our research highlights how specific microbial taxa contribute to nutrient cycling and the decomposition of organic materials, processes crucial for maintaining water quality. By identifying key microbial taxa that are involved in these processes, management strategies can be tailored to support or enhance these natural functions, potentially leading to more effective interventions for controlling eutrophication. For example, strategies could focus on reducing nutrient loads in areas where key microbes are active in nutrient cycling, thus mitigating the risk of harmful algal blooms. The study by Herren et al. [36] underscores the role of keystone taxa in stabilizing microbial communities, which suggests that maintaining or enhancing the populations of such taxa could play a significant role in sustaining water quality. This research demonstrates that small subsets of highly connected keystone taxa can predict whole-community compositional changes, which is crucial for managing ecosystem responses to environmental fluctuations [36].

Overall, understanding these adaptive mechanisms can help predict potential shifts in microbial dynamics in response to warming temperatures, altered precipitation patterns, or other climatic changes. This predictive capacity is crucial for developing adaptive management strategies that can accommodate future changes, ensuring the long-term health and stability of Lake Mendota and other freshwater ecosystems. As we continue to face global environmental challenges, such research is vital for developing strategies to preserve the health and stability of aquatic ecosystems.

## 4 Materials and Methods

### 4.1 Data description and data processing

#### 4.1.1 Data description

This study utilizes two distinct datasets derived from bulk metagenomic and metatranscriptomic sequencing, focusing on the microbial ecology of Lake Mendota located in Wisconsin, USA [15]. In summary, samples were collected during the open-water period from Lake Mendota, a dimictic, seasonally anoxic lake. The samples were collected at different depths, weekly, over the course of a season, therefore capturing a range of environmental conditions. For each sample, DNA was extracted for metagenomic sequencing, and RNA was extracted for metatranscriptomic sequencing. This allowed us to analyze both the genetic potential (bulk-DNA) and the actual gene expression (bulk-RNA) of the microbial communities. The metagenomes were binned into MAG, each representing a distinct microbe (and its genome), and metatranscriptomic reads were mapped onto the MAGs to determine their activity over time and space. The metagenomic dataset comprises raw counts of metagenomic reads aligned to 431 MAGs, representing a species within the Bacteria and Archaea domains of life. These MAGs were cataloged across 16 distinct environmental samples, chosen to capture a wide range of environmental conditions such as variations in depth, temperature, and nutrient availability. This diverse sampling provides a rich matrix for analyzing microbial diversity and its spatial and temporal fluctuations within the lake. Metagenomics reveals the “potential” metabolic activity of microbes, whereas metatranscriptomics determines their “actual” activity in the environment. Since the goal of our study is to identify key taxa in the microbial network, the metatranscriptomic analysis further validates the “key taxa” findings. Therefore, the metatranscriptomic dataset includes raw counts of RNA reads aligned to the same set of 431 MAGs, offering insights into the active gene expression profiles of these microbial communities. This dataset allows for an in-depth examination of the functional dynamics and metabolic activities of the microbes in response to environmental conditions, complementing the genomic data with a layer of functional information.

#### 4.1.2 Sampling Information

The samples were collected from Lake Mendota, a well-studied freshwater lake known for its dynamic ecological processes. Sampling was conducted over several months, capturing seasonal variations that influence microbial communities. Each sample was taken at specific depths and locations to ensure a representative understanding of both horizontal and vertical microbial distribution. The dataset associated with this research encompasses several environmental parameters meticulously recorded during the sample collection from Lake Mendota. Each parameter is stored in specific columns within the dataset, including temperature, dissolved oxygen, and pH levels. For details, see Table S1 in the Supplementary Material.

#### 4.1.3 Data Selection Criteria and Normalization

For the analysis within this study, specific filtering steps were applied to enhance the dataset’s integrity and relevance: The column “do_sat (%)”, which represents dissolved oxygen saturation in percentage, was excluded from this analysis. This decision is based on its redundancy with the do_raw (mg/L) column, which provides a direct and precise measurement of dissolved oxygen in milligrams per liter. By focusing on the “do_raw data”, we ensure that the dataset remains streamlined and retains only the most analytically useful information, thereby enhancing the precision of ecological assessments.

Normalization is another critical step in metagenomic data analysis to adjust for variations in sequencing depth and sampling disparities. Two different normalization methods were applied across various sections of our analysis: 1-Row sum normalization [37]: In the initial data processing phase, the row sum normalization method was applied. This method standardizes the data by dividing the counts in each sample by the total counts of that sample. This normalization technique is essential for adjusting for different sequencing depths across samples, allowing for comparative analysis across the dataset. It is particularly effective in highlighting relative differences in microbial population structures between samples, thus facilitating a better understanding of microbial ecology. 2-Min-Max normalization [38]: During the analysis involving `CARLasso`, Min-Max normalization was utilized. This technique scales each feature to a fixed range, typically between 0 and 1, according to the 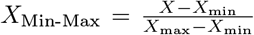 formula. Where, *X* denotes the original value, *X*_min_ is the minimum value of the feature, and *X*_max_ is the maximum value. Min-max normalization is crucial for regression analyses in `CARlasso` models as it ensures that each environmental feature contributes equally to the model, avoiding biases due to varying scales of measurement. Normalization improves the accuracy and reliability of the analysis by ensuring consistent scaling across all features.

### 4.2 Feature Selection

#### 4.2.1 Network Construction using SPIEC-EASI

In our study, we utilized the `SPIEC-EASI` algorithm to construct a microbiome interaction network. This method infers conditional dependencies among MAGs by calculating the inverse covariance matrix, also known as the precision matrix. This matrix highlights direct relationships between variables while controlling for others. A non-zero value in this matrix between two variables indicates a direct relationship. Using the precision matrix, we constructed a network representing the conditional dependencies among MAGs. This network outlines the intricate interdependencies within the microbial community and serves as the foundation for further feature selection processes. We define key taxa as MAGs that are highly connected and influential. A highly connected MAG interacts with many other MAGs, while an influential MAG significantly impacts the microbial community’s structure and function, such as playing a crucial role in nutrient cycling or affecting the growth and survival of other microorganisms. Feature selection was performed on the constructed network by calculating the connectivity, or degree, of each MAG. The degree indicates the number of direct connections a node (MAG) has with other nodes. We selected the top 15 MAGs based on their degree values, considering them as key players within the microbial community due to their high connectivity. This selection criterion highlights the most influential MAGs in terms of network structure and ecological impact for further analysis.

#### 4.2.2 Taxonomic Classification

The Genome Taxonomy Database Toolkit (GTDB-Tk) provides objective taxonomic assignments for bacterial and archaeal genomes based on the GTDB. GTDB-Tk is computationally efficient and able to classify thousands of draft genomes in parallel [39]. GTDB-tk was previously used to classify each MAG [15]. To facilitate further analysis, we used a custom R script to match the MAG names with their corresponding taxonomic classifications. This step enables us to provide ecological context for the network findings. This enables use to provide ecological context for the network findings. This methodological approach, combining `SPIEC-EASI` with degree-based feature selection, provides a robust framework for elucidating the structural and functional dynamics of microbial communities.

### 4.3 Model descriptions: Bayesian sparse microbial network via a Gaussian chain graph (CARlasso)

In our study, we employed the `CARlasso` method [23, 24] within a Bayesian framework to construct a Gaussian chain graph. This method is implemented using the `CARlasso` package in R. `CARlasso` is uniquely suited for ecological and biological data analysis. It allows for the inference of chain graphs by integrating both microbial abundances and environmental or experimental conditions as nodes within the graph. The Gaussian chain graph constructed using `CARlasso` distinguishes between two types of nodes: 1-Correlated Responses: These nodes represent the microbial abundances. They are essential for understanding the dynamics within the microbial community and how these entities interact with each other under various conditions. 2-Predictors: These nodes account for environmental or experimental conditions that affect the microbial responses. This inclusion allows the model to capture the impact of external factors on microbial behavior and interactions. Utilizing the key MAGs identified through our network analysis, along with all measured environmental parameters, we constructed a comprehensive network using the `CARlasso` model. This network serves multiple purposes: 1-Integrative Analysis: By incorporating both MAGs and environmental parameters, the network offers a holistic view of the microbial ecosystem, showcasing how environmental conditions directly affect microbial abundance and interactions. 2-Enhanced Interpretability: The model facilitates a deeper understanding of the ecological roles of selected MAGs and their interactions with environmental factors, providing valuable insights into ecosystem functioning and stability. Using `CARlasso` to construct a Gaussian chain graph enhances our understanding of microbial community dynamics and aligns with biological expectations of conditional interactions. This approach proves to be a robust tool for ecological network analysis.

## Code and Data availability

Data and all reproducible scripts are available in the repository at https://github.com/solislemuslab/lake-microbiome-data-analysis. For further inquiries regarding the data, please contact Karthik Anan-tharaman at.

## Acknowledgements

We are grateful to the Center for Limnology and the Long-Term Ecological Research – North Temperate Lakes group for supporting field sampling. PQT was funded by the Natural Science and Engineering Research Council of Canada (NSERC) doctoral fellowship, the Anna Grant Birge Memorial Award from the Center for Limnology, and the Baldwin Distinguished Graduate Fellowship from the Department of Bacteriology at the University of Wisconsin-Madison. DNA sequencing was conducted at the DOE Joint Genome Institute, a DOE Office of Science User Facility, via a Community Science Program New Investigator award to KA and PQT (award number 506328). This work was supported by the National Science Foundation [DEB-2144367 to CSL, and DBI-2047598 to KA] and the USDA National Institute of Food and Agriculture (NIFA) under grant: Hatch 1025641.

## Author Contributions

CSL and KA worked in Conceptualization, formulating initial ideas and defining research goals. PQT managed Data Curation, preparing data for analysis and future use. QY led the Formal Analysis, applying statistical techniques to analyze the study data and taking the lead in Writing – the original draft, and preparing the initial manuscript. RA contributed to the Formal analysis by assisting with statistical methods and model fitting. QY, RA, PQT, and CSL played key roles in interpreting the results and Writing – review & editing, contributing to the critical review and refinement of the manuscript. All authors participated in Validation, ensuring the findings’ reproducibility and integrity, and approved the final manuscript, affirming their shared responsibility.

## SUPPLEMENTARY MATERIAL

**Table S1:**
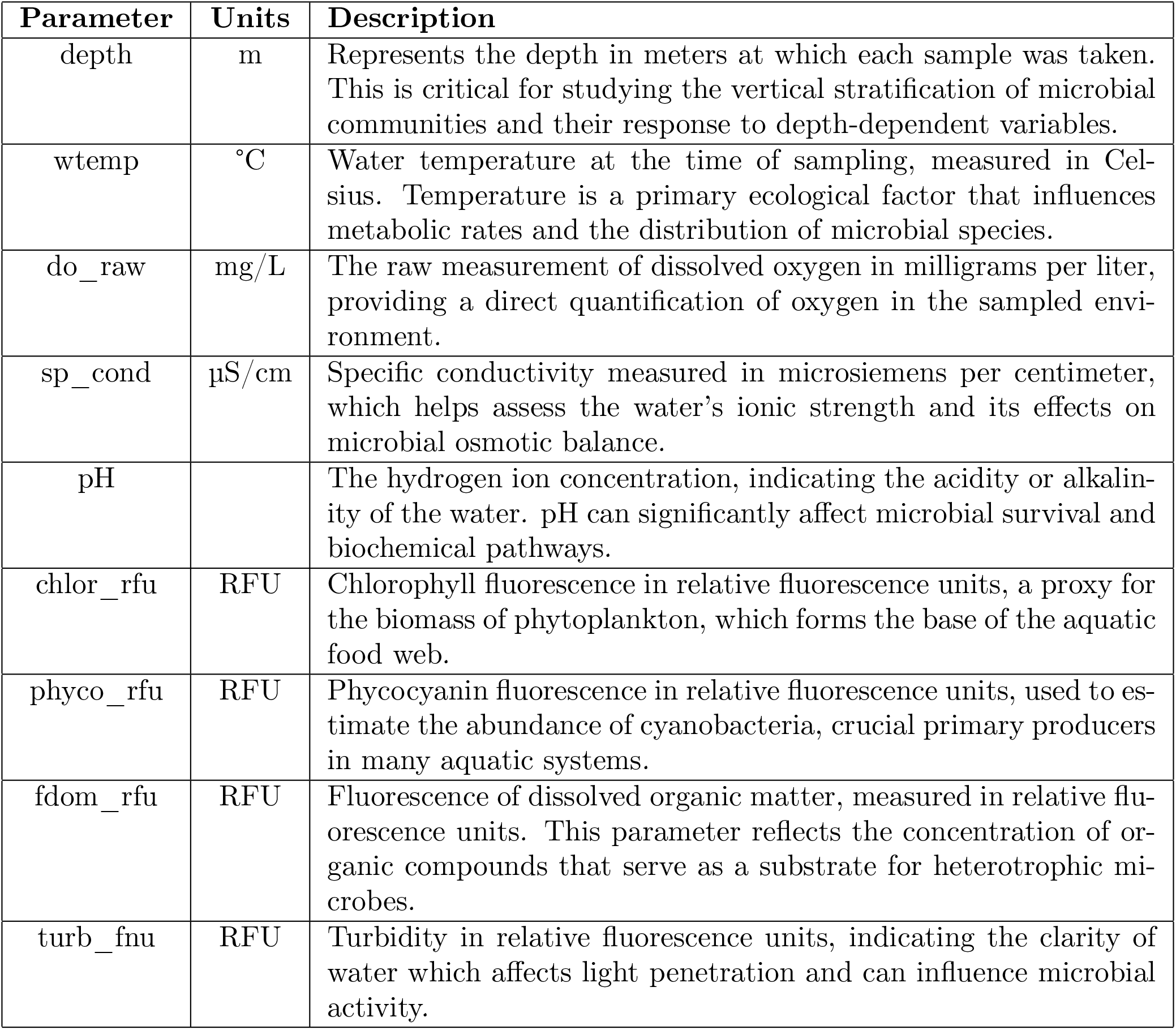
Descriptions of environmental parameters used in the study.

**Figure S1:**
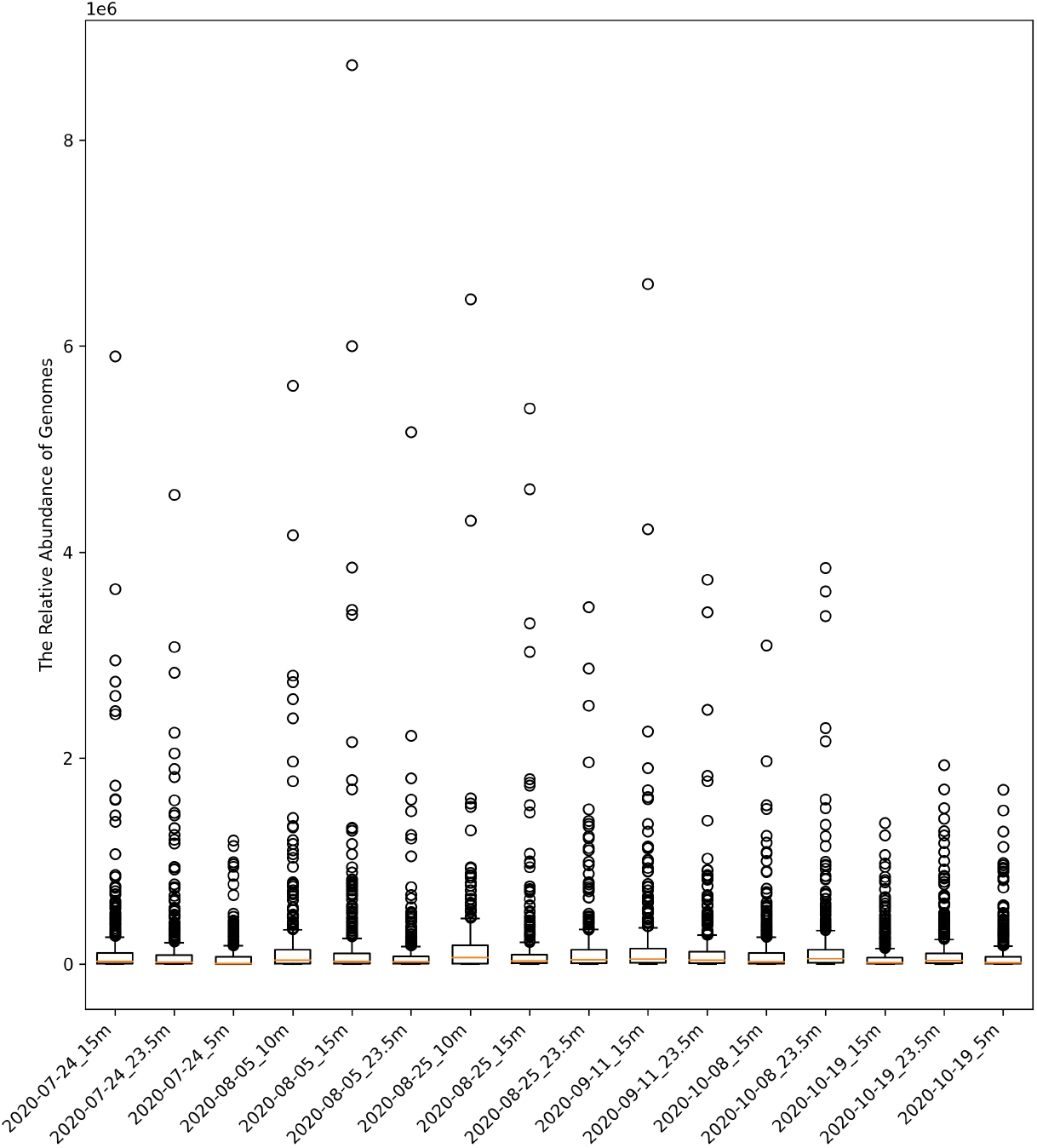
Boxplot for distribution of the relative abundance of genome per sample after using using row sum normalization method. The x-axis labels represent the sampling dates and depths. Each label consists of the year, month, day, and the depth at which the sample was taken. For example, the label “2020-07-24_15m” indicates that the sample was collected on July 24, 2020, at a depth of 15 meters.

**Figure S2:**
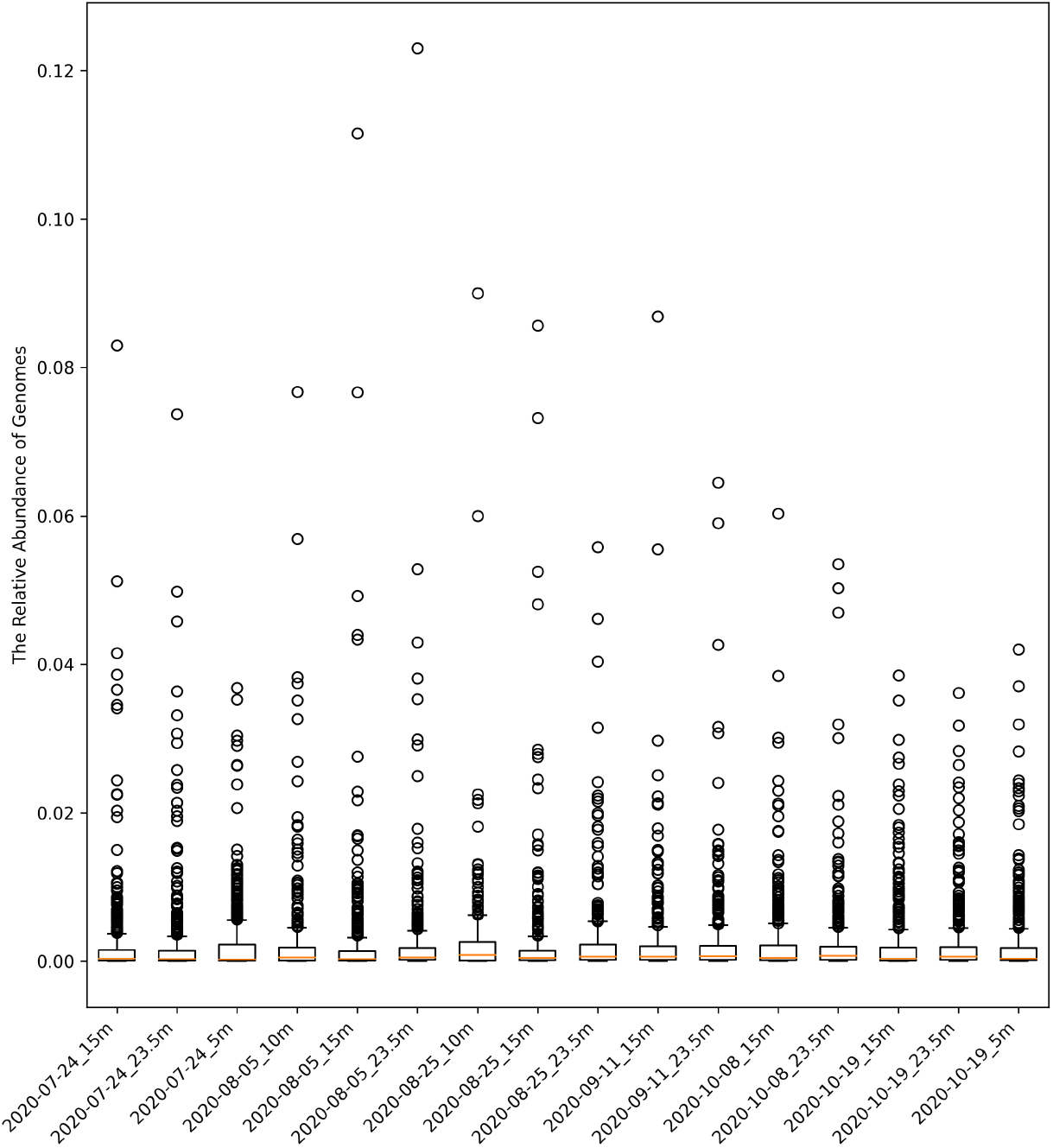
Boxplot for distribution of the relative abundance of genome per sample after using using row sum normalization method. The x-axis labels represent the sampling dates and depths. Each label consists of the year, month, day, and the depth at which the sample was taken. For example, the label “2020-07-24_15m” indicates that the sample was collected on July 24, 2020, at a depth of 15 meters.

**Figure S3:**
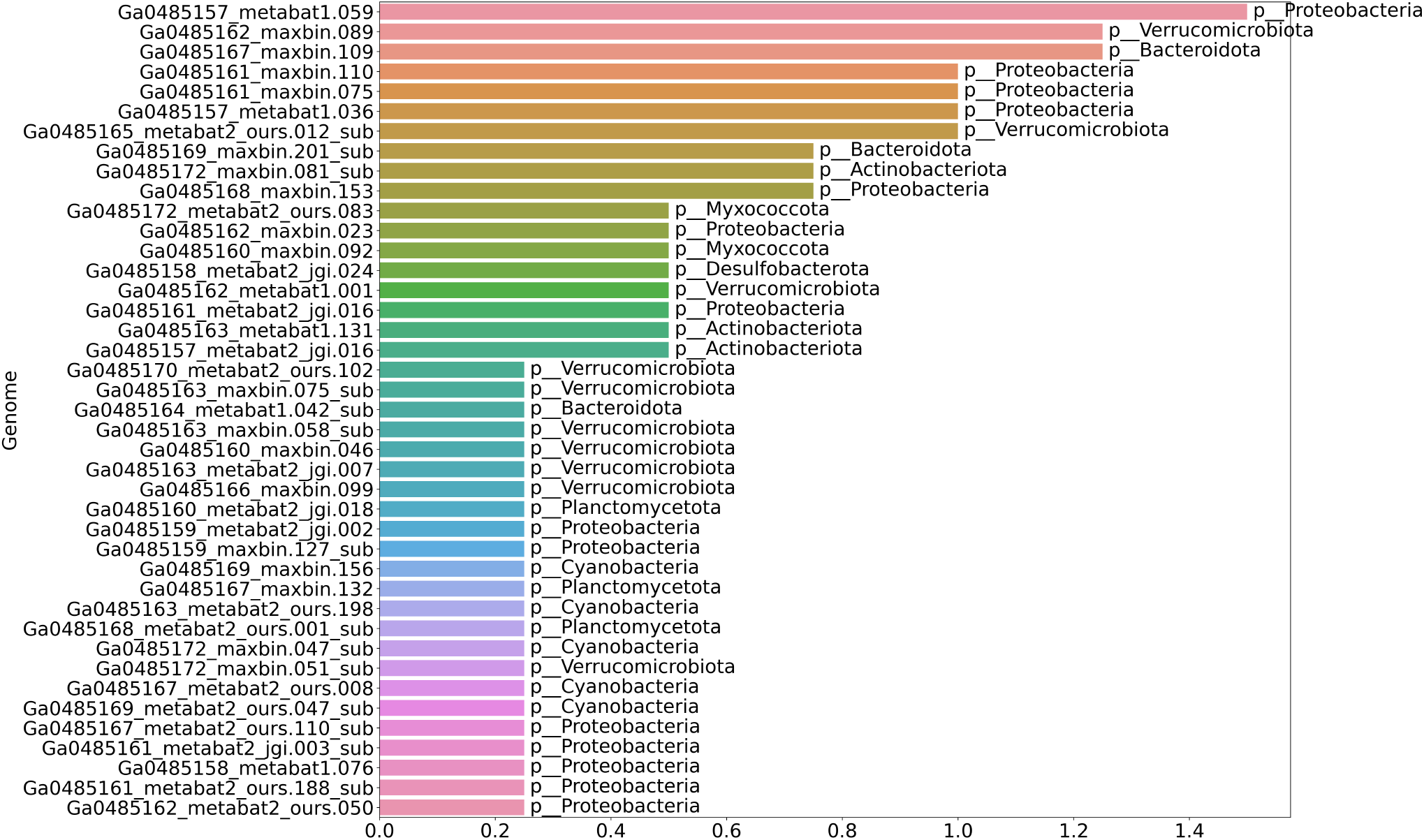
Cumulative ratio of the appearance of the most prevalent genomes across different time periods and depths. The x-axis represents the cumulative ratio of each genome, while the y-axis lists the genomes. Each bar indicates the cumulative appearance ratio of specific genomes for given months and depths, with added labels indicating the phylum to which each genome belongs. Notably, genomes like MAG Ga0485157_metabat1.059, classified under the phylum Proteobacteria, show high prevalence, suggesting their significant roles in the microbial community structure.

**Figure S4:**
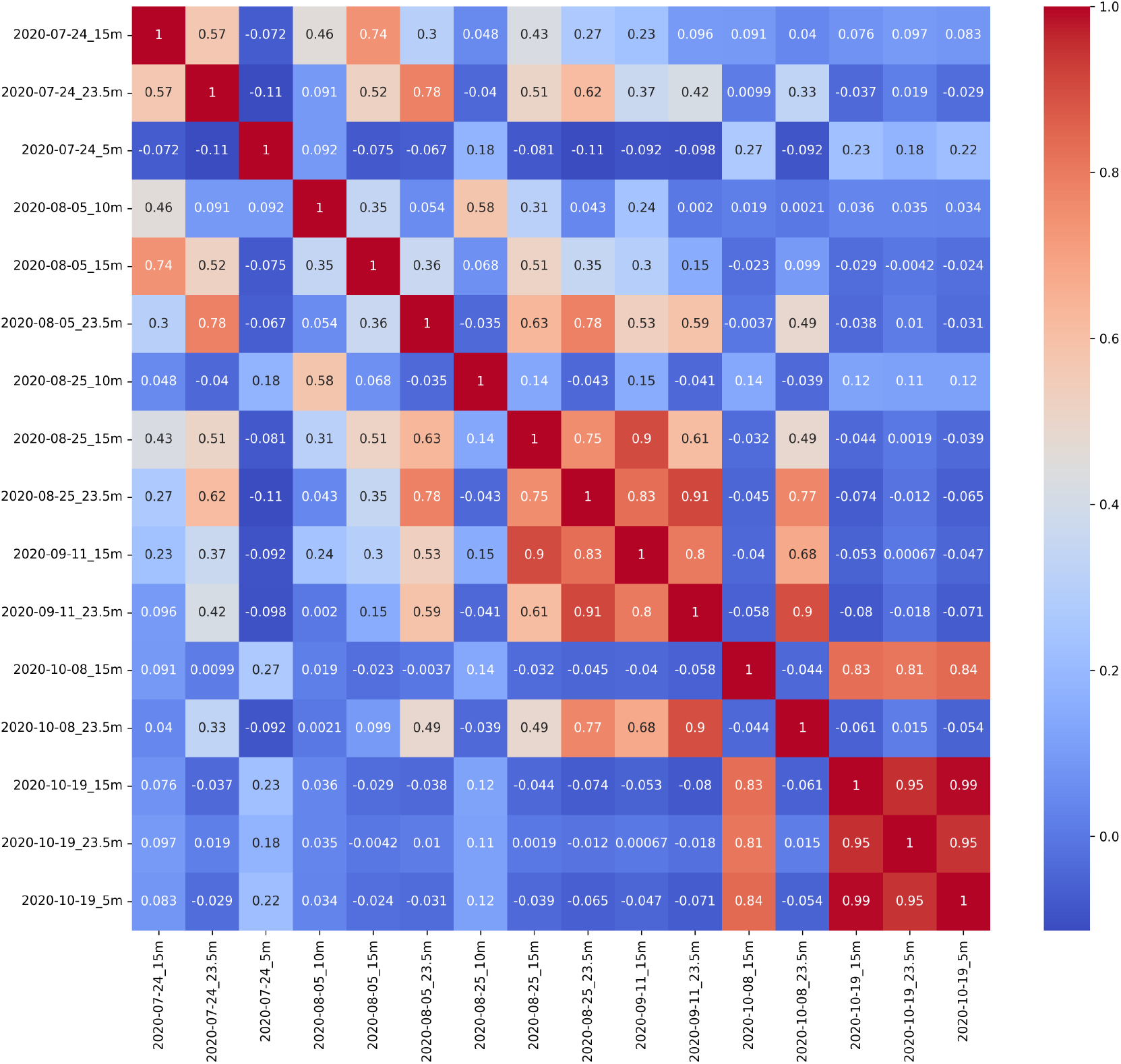
Pearson’s correlation heatmap displaying linear relationships among microbial samples. Notable strong correlations are observed between samples collected from different depths on the same date, enhancing our understanding of the microbial community structure.

**Figure S5:**
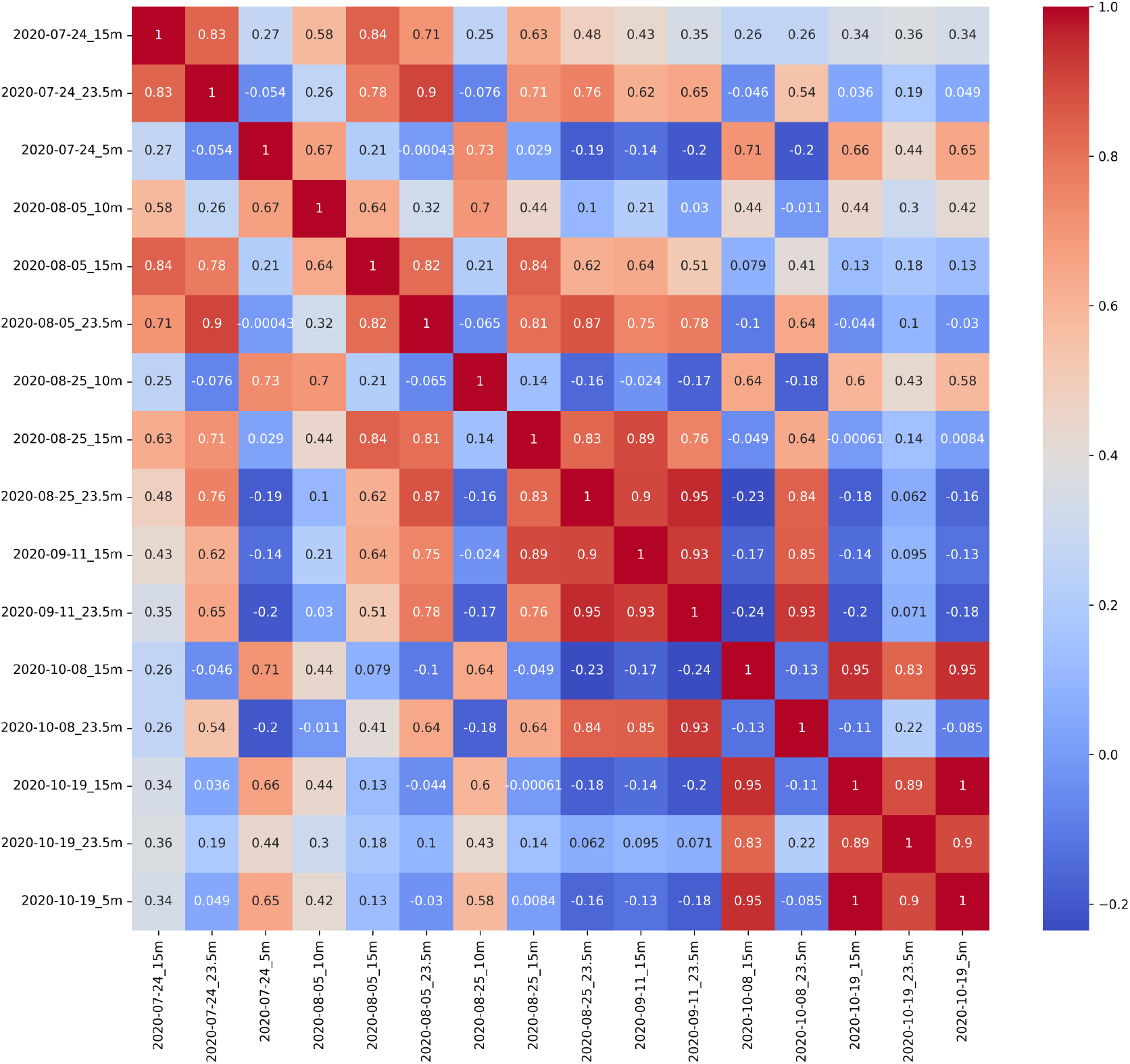
Spearman’s correlation heatmap showing monotonic relationships among the samples. This analysis highlights both linear and non-linear associations, providing a comprehensive view of the microbial interactions within the ecosystem.

## Notes

https://github.com/solislemuslab/lake-microbiome-data-analysis

### Competing Interest Statement

The authors have declared no competing interest.

