## supplemental file for "Unraveling Keystone Taxa: Interactions Within Microbial Networks and Environmental Dynamics in Lake Mendota"

November 11, 2024

| **Parameter** | **Units** | **Description** |
|---|---|---|
| depth | m | Represents the depth in meters at which each sample was taken. This is critical for studying the vertical stratification of microbial communities and their response to depth-dependent variables. |
| wtemp | ˚C | Water temperature at the time of sampling, measured in Celsius. Temperature is a primary ecological factor that influences metabolic rates and the distribution of microbial species. |
| do_raw | mg/L | The raw measurement of dissolved oxygen in milligrams per liter, providing a direct quantification of oxygen in the sampled environment. |
| sp_cond | µS/cm | Specific conductivity measured in microsiemens per centimeter, which helps assess the water's ionic strength and its effects on microbial osmotic balance. |
| pH | | The hydrogen ion concentration, indicating the acidity or alkalinity of the water. pH can significantly affect microbial survival and biochemical pathways. |
| chlor_rfu | RFU | Chlorophyll fluorescence in relative fluorescence units, a proxy for the biomass of phytoplankton, which forms the base of the aquatic food web. |
| phyco_rfu | RFU | Phycocyanin fluorescence in relative fluorescence units, used to estimate the abundance of cyanobacteria, crucial primary producers in many aquatic systems. |
| fdom_rfu | RFU | Fluorescence of dissolved organic matter, measured in relative fluorescence units. This parameter reflects the concentration of organic compounds that serve as a substrate for heterotrophic microbes. |
| turb_fnu | RFU | Turbidity in relative fluorescence units, indicating the clarity of water which affects light penetration and can influence microbial activity. |

Table 1: Descriptions of environmental parameters used in the study.

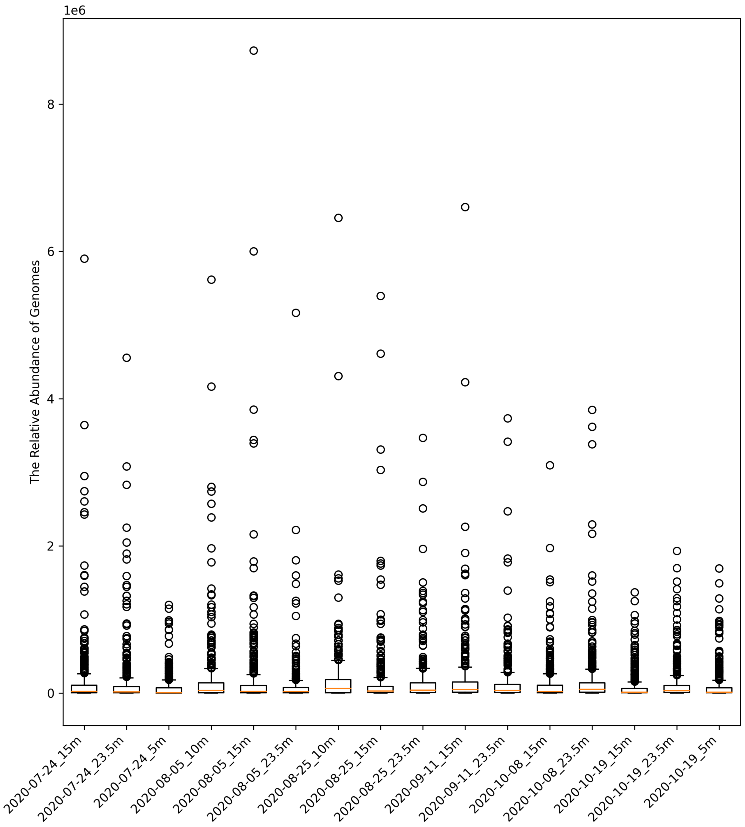

Figure 1: Boxplot for distribution of the relative abundance of genome per sample before normalization. The x-axis labels represent the sampling dates and depths. Each label consists of the year, month, day, and the depth at which the sample was taken. For example, the label “2020-07-24_15m" indicates that the sample was collected on July 24, 2020, at a depth of 15 meters.

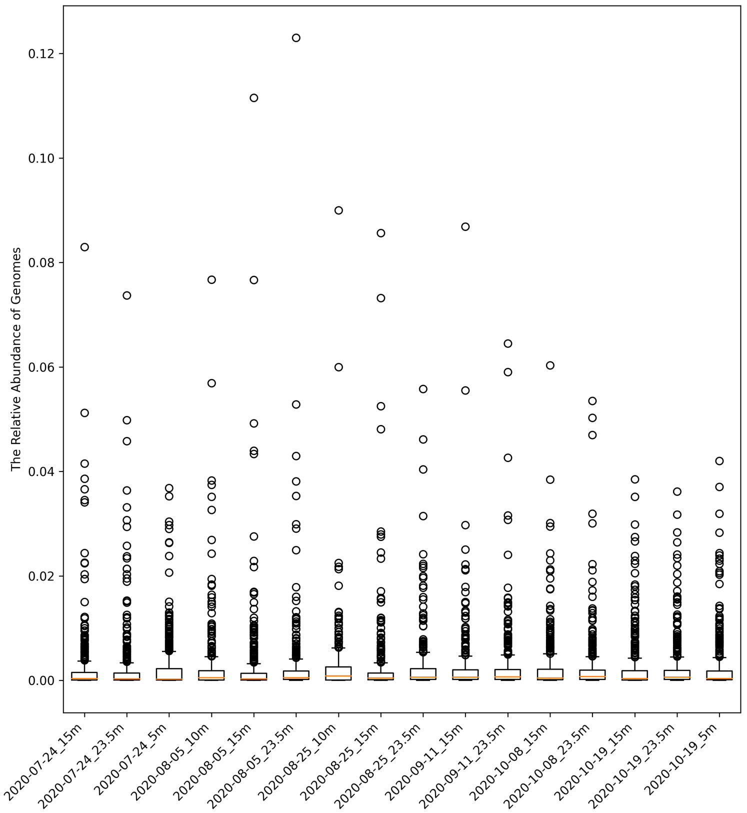

Figure 2: Boxplot for distribution of the relative abundance of genome per sample after using using row sum normalization method. The x-axis labels represent the sampling dates and depths. Each label consists of the year, month, day, and the depth at which the sample was taken. For example, the label "2020-07-24_15m" indicates that the sample was collected on July 24, 2020, at a depth of 15 meters.

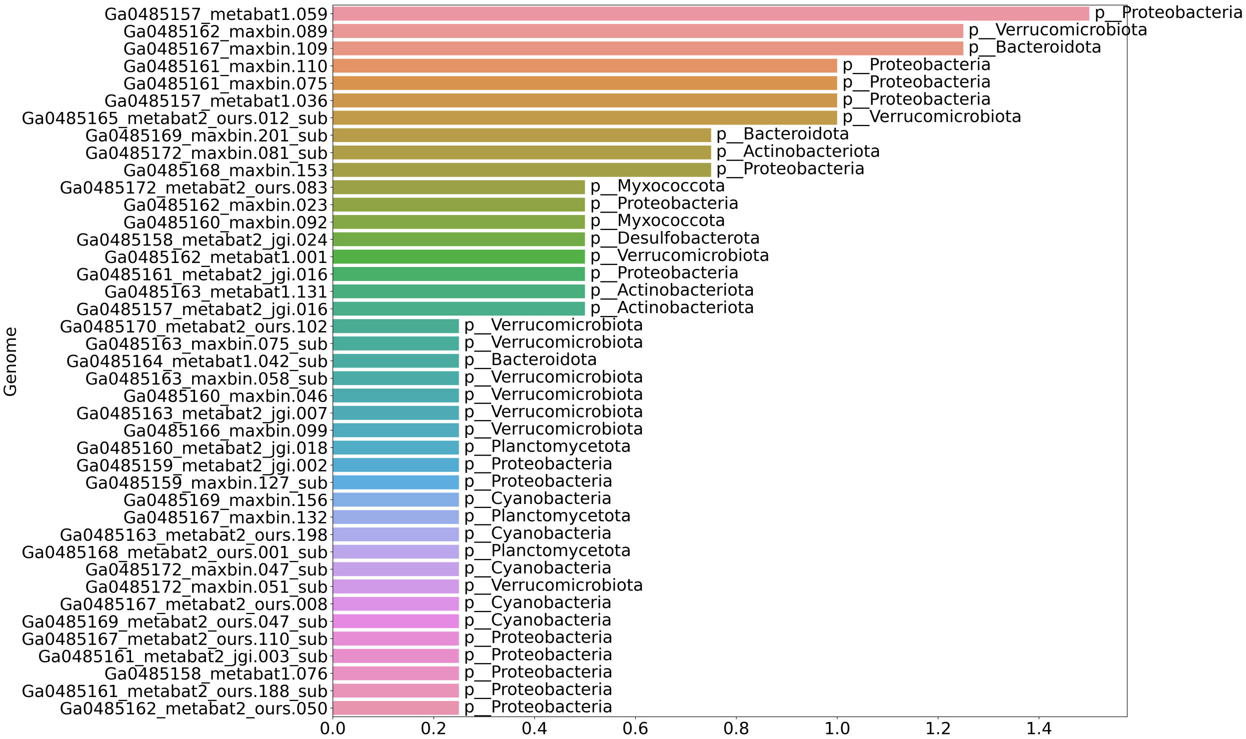

Figure 3: Cumulative ratio of the appearance of the most prevalent genomes across different time periods and depths. The x-axis represents the cumulative ratio of each genome, while the y-axis lists the genomes. Each bar indicates the cumulative appearance ratio of specific genomes for given months and depths, with added labels indicating the phylum to which each genome belongs. Notably, genomes like MAG Ga0485157_metabat1.059, classified under the phylum Proteobacteria, show high prevalence, suggesting their significant roles in the microbial community structure.

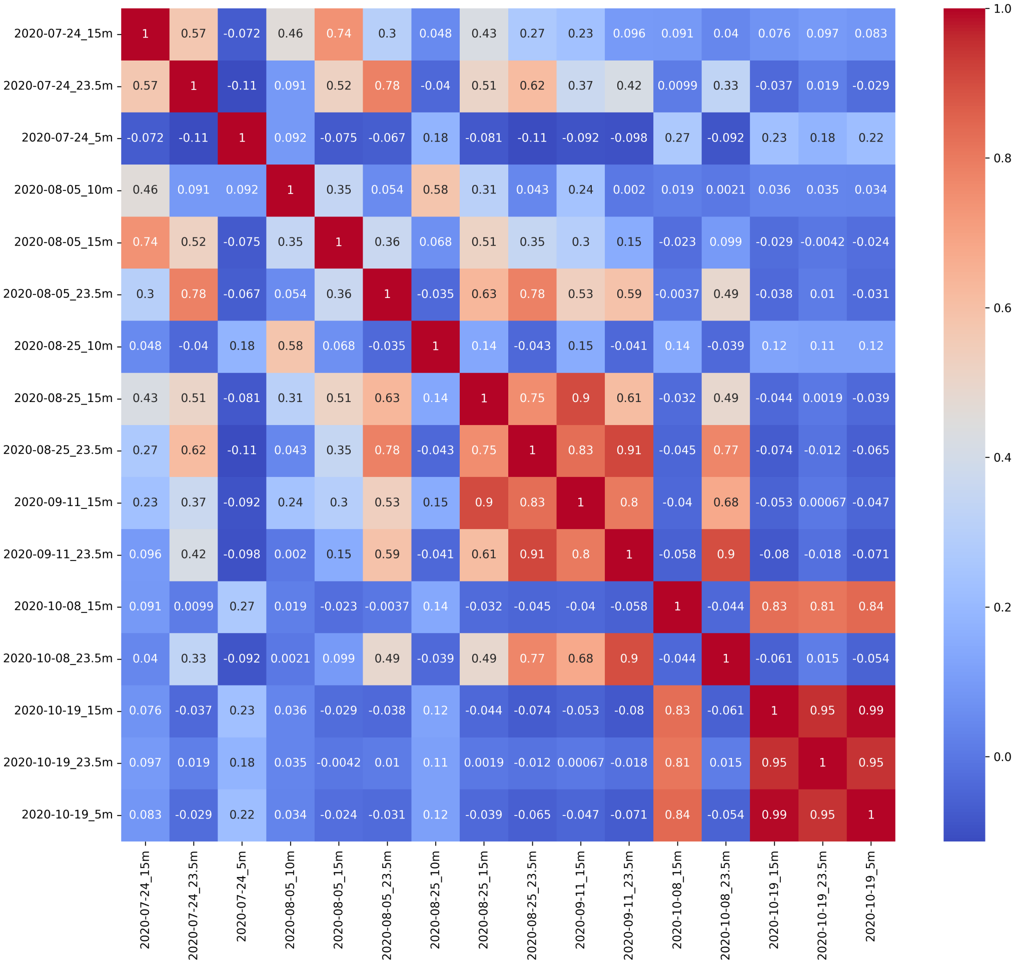

Figure 4: Pearson's correlation heatmap displaying linear relationships among microbial samples. Notable strong correlations are observed between samples collected from different depths on the same date, enhancing our understanding of the microbial community structure.

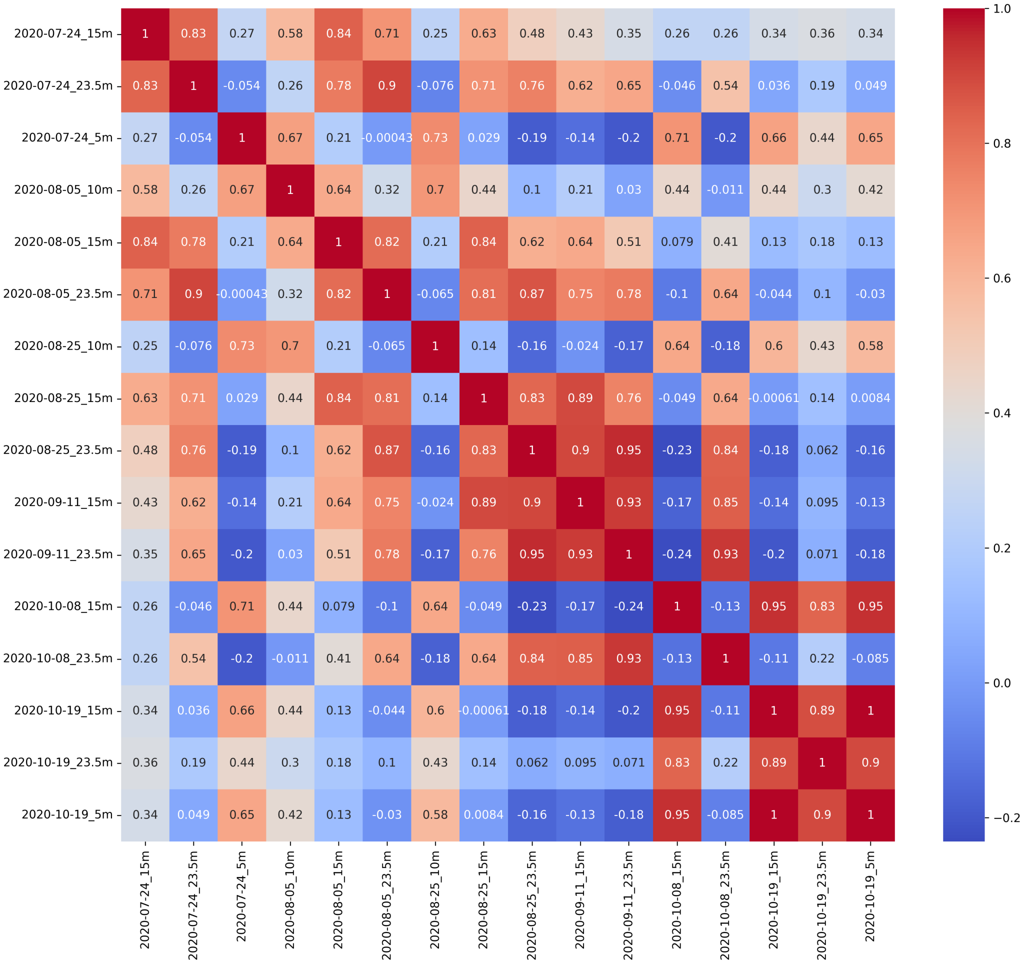

Figure 5: Spearman's correlation heatmap showing monotonic relationships among the samples. This analysis highlights both linear and non-linear associations, providing a comprehensive view of the microbial interactions within the ecosystem.
